# Video-based biomechanical analysis captures disease-specific movement signatures of different neuromuscular diseases

**DOI:** 10.1101/2024.09.26.613967

**Authors:** Parker S. Ruth, Scott D. Uhlrich, Constance de Monts, Antoine Falisse, Julie Muccini, Sydney Covitz, Shelby Vogt-Domke, John Day, Tina Duong, Scott L. Delp

**Affiliations:** Department of Computer Science, Stanford University; Department of Mechanical Engineering, University of Utah; Department of Neurology, Stanford Medicine; Department of Bioengineering, Stanford University; Department of Radiology, Stanford Medicine; Department of Mechanical Engineering, Stanford University; Department of Orthopaedic Surgery, Stanford Medicine

## Abstract

**Background:** Assessing human movement is essential for diagnosing and monitoring movement-related conditions like neuromuscular disorders. Timed function tests (TFTs) are among the most widespread assessments due to their speed and simplicity, but they cannot capture disease-specific movement patterns. Conversely, biomechanical analysis can produce sensitive disease-specific biomarkers but is traditionally confined to laboratory settings. Recent advances in smartphone video-based biomechanical analysis enable quantification of 3D movement with the ease and speed required for clinical settings. However, the potential of this technology to offer more sensitive assessments of human function than TFTs remains untested.

**Methods:** To compare video-based analysis against TFTs, we collected an observational dataset from 129 individuals: 28 with facioscapulohumeral muscular dystrophy, 58 with myotonic dystrophy, and 43 controls with no diagnosed neuromuscular condition. We used OpenCap, a free open-source software tool, to capture smartphone video-based biomechanics of nine different movements in a median time of 16 minutes per participant. From these recordings we extracted 34 interpretable movement features. Using these features, we evaluated the ability of video-based biomechanics to reproduce four TFTs (10-meter walk, 10-meter run, timed up-and-go, and 5-time sit-to-stand) while capturing additional disease-specific signatures of movement.

**Results:** Video-based biomechanical analysis reproduced all four TFTs (r > 0.98) with similar test-retest reliability. In addition, video metrics outperformed TFTs at disease classification (p = 0.021). Unlike TFTs, video-based biomechanical analysis identified disease-specific signatures of movement such as differences in gait kinematics that are not evident in TFTs.

**Conclusion:** Video-based biomechanical analysis can complement existing functional movement assessments by capturing more sensitive, disease-specific outcomes from human movement. This technology enables digital health solutions for assessing and monitoring motor function, complementing traditional clinical outcome measures to enhance care, management, and clinical trial design for movement-related conditions.

**Description:** This study demonstrates that smartphone video-based biomechanical analysis can accurately replicate traditional timed function tests (TFTs) — commonly used to diagnose and assess movement-related conditions like neuromuscular disorders — while also capturing disease-specific movement patterns that TFTs fail to detect. By enabling sensitive and interpretable assessments in clinical settings, this approach offers a scalable and objective tool to enhance diagnosis, monitoring, and clinical trial design for neuromuscular disorders.

## Introduction

Quantitative measurements of human movement can improve diagnosis, monitoring, and understanding of movement-related disorders. The state of the art for measuring human movement involves marker-based motion capture and force plates, which cost hundreds of thousands of dollars and require hours of expert labor.^1^ While laboratory-based methods are the gold standard,^2^ real-world clinical applications require greater speed and convenience.

Timed function tests (TFTs), which assess gross motor function by measuring the time taken to complete an activity, have wide clinical adoption due to their speed and simplicity. However, TFTs are less informative than laboratory-based biomechanical analysis. Reducing human movement to a single time score overlooks movement quality factors like compensatory gait changes or limited joint range-of-motion, which may better reflect disease progression. Thus, there is a gap between the rich measurement tools used in biomechanics research and those deployed in clinical settings.

With the ongoing development of disease-modifying therapies for neuromuscular diseases, improved measures of mobility and function are crucial for evaluating treatment efficacy, clinical decision-making, and disease management.^3,4^ The sensitivity and accuracy of clinical endpoints greatly impact trial design and therapy advancement. For slower-progressing diseases, TFTs lack sensitivity to subtle changes in muscle function. Coarse metrics can fail to detect natural progression and the effects of therapeutic interventions. For example, individuals with early-stage disease may retain normal walking speeds (and hence normal TFT times) but display compensations or decreased movement quality indicating impaired muscle function. There is a strong clinical need for functional assessments sensitive to the unique phenotypical signatures of muscle dysfunction across neuromuscular diseases.^5^ Prior work has demonstrated the promise of digital movement biomarkers to better quantify disease progression in several neuromuscular diseases, including Duchenne muscular dystrophy,^6^ Friedreich’s ataxia,^7^ and Parkinson’s disease.^8^ Using full-body inertial measurement unit sensors and laboratory-based motion capture, these studies show how movement biomarkers can better predict disease trajectory and are more sensitive to short-term progression than standard clinical tests.

Here, we focus on two slow-progressing neuromuscular diseases: facioscapulohumeral muscular dystrophy (FSHD) and myotonic dystrophy (DM). There are no approved digital outcome measures for these conditions, and their slow progression necessitates sensitive outcome measures to accelerate the development of treatments. Both conditions impact gross motor function (e.g., reduce walking speed), but they affect different muscle groups, leading to distinct movement signatures. FSHD causes weakness in facial, scapulothoracic, shoulder, trunk, and ankle dorsiflexor muscles, resulting in impaired locomotion and overhead reaching.^9,10^ DM primarily causes weakness in the lower trunk and distal muscles of the ankle, hand, trunk, knee, and hip, resulting in difficulty rising from a chair, squatting, and activities of daily living.^11^

Quantifying human movement with video can provide richer metrics with minimal added burden.^12^ For example, machine learning models can predict clinical scores of Parkinson’s disease severity from videos of gait^13^ and sit-to-stand transitions.^14^ The speed and ease of video capture facilitates scalable data collection in clinical and remote settings. For example, at-home videos of sit-to-stand transitions predict osteoarthritis and health outcomes.^15^ Video capture is rich and flexible enough to tailor metrics for individual diseases. For example, “reachable workspace” is a video-derived metric quantifying upper-body functional volume developed for FSHD.^16^ Reachable workspace is validated against other upper-limb metrics,^17^ reflects longitudinal progression,^18^ correlates with patient reported-outcomes, and is used as an outcome measure in clinical trials.^19^

We previously developed OpenCap, an open-source tool that computes three-dimensional (3D) human kinematics (i.e., joint positions and angles) from smartphone videos.^20^ By eliminating time-consuming analysis and specialized equipment, this greatly expands the scale and accessibility of human movement analysis. For example, OpenCap can detect subtle movement patterns — such as steppage gait or circumduction gait — that go beyond simple metrics like walking speed.^21^

This study aimed to compare video-based 3D biomechanical analysis using OpenCap (Figure 1) to TFTs for quantifying movement of individuals with FSHD and DM. Specifically, we investigated two research questions: (1) Can OpenCap video-based metrics accurately reproduce clinician-graded TFTs in individuals with and without neuromuscular diseases? (2) Can OpenCap video-based biomechanical assessment capture disease-specific movement patterns in FSHD and DM that are more informative than TFT times alone? To answer these questions, we collected a dataset of OpenCap recordings and TFTs, extracted video-based movement features, and explored the potential for video-based digital metrics of human movement to both reproduce TFTs and provide additional relevant movement information about two neuromuscular diseases.

**Figure 1:**
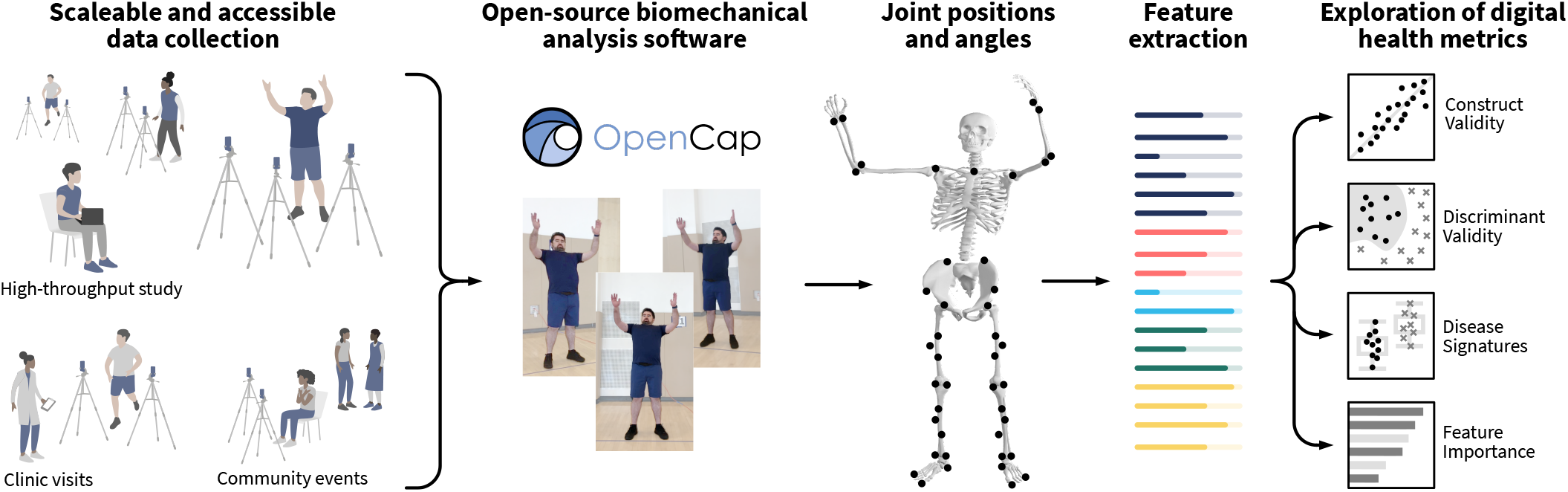
Video-based biomechanical analysis. Video-based biomechanical analysis enables scalable and portable data collection for studying human movement. We used the open-source OpenCap application**^20^**to convert smartphone videos to three-dimensional anatomical marker positions and joint angles. Leveraging clinical domain knowledge, we engineered a set of video-based metrics describing aspects of human motion known to have salience for the neuromuscular diseases we study. These features are then used to explore the design of novel digital health metrics. (Identifiable photos shared with informed consent.) Source: created by authors

## Methods

### Participant screening and recruitment

One hundred twenty-nine individuals completed the study (Table 1), including 28 with FSHD, 58 with DM (49 myotonic dystrophy type 1 [DM1], 9 myotonic dystrophy type 2 [DM2]), and 43 controls with no diagnosed neuromuscular condition. (For detailed participant demographics see Table S1.) Prior to participation, participants provided written informed consent to a protocol approved by the Stanford University Institutional Review Board (IRB IRB00005136). The OpenCap system is HIPAA compliant and has been approved by the Stanford University Offices of Privacy and Data Security to process and store identifiable personal health information. Participants provided consent to the collection, processing, and optional sharing of their identifiable video data.

**Table 1:**
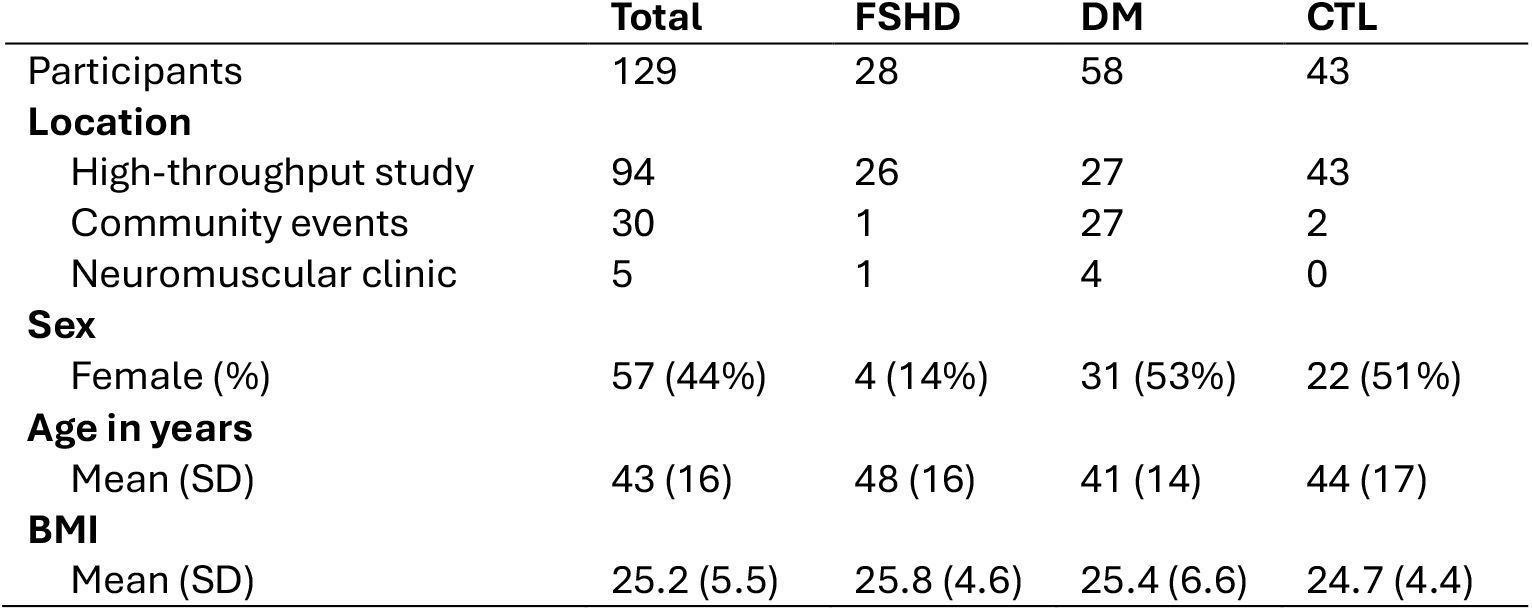
Study participant characteristics. (FSHD = facioscapulohumeral muscular dystrophy, DM = myotonic dystrophy, CTL = control)

Inclusion criteria were the ability to rise from a chair and ambulate independently. Participants in the control group had no neuromuscular condition, movement disorder, or injury affecting movement in the last six months. Participants with movement disorders had confirmed diagnosis of DM (DM1 or DM2) or FSHD. Participants were recruited in three settings: 94 individuals participated in a high-throughput data collection, primarily in a university gymnasium using multiple parallel OpenCap motion capture stations; 30 participants were recruited on-site at support group meetings and national conferences; 5 participants were recruited during clinic visits.

### Selection of physical activities

In consultation with the clinical domain experts on the study team, we selected a set of nine upper- and lower-body movements (Table S2) representing isolated muscle groups and whole-body function. We chose run, walk, timed up-and-go, and sit-to-stand movements because they are common TFTs. We added a 30-second calf raise to test balance and plantarflexor strength and a jump task to test maximal force of lower-extremity extensors. Finally, we chose three upper-body movements: shoulder abduction, elbow flexion, and arm range-of-motion. The arm abduction task is used in the existing Brooke Upper Extremity Scale, which ordinally scores functional ability of overhead arm abduction, hand-to-mouth movement, and distal hand function.^22,23^ Together, these activities represent a wide spectrum of functional movement amenable to video-based biomechanical analysis.

### Domain expert-informed feature engineering

We leveraged the expertise of clinical research team members and previously reported movement patterns in FSHD and DM to select a set of 34 interpretable, activity-specific features from the kinematic data (Table S3). To design features, we discussed common movement patterns exhibited by individuals with DM and FSHD and examined representative videos and skeletal kinematics from the dataset. The clinicians verbally identified common movement impairments and compensations, and engineers designing algorithms to quantify clinical observations. Some participants were unable to complete ankle-based activities (calf raise and jump); these missing values were imputed with zeros.

### Scalable video-based human movement assessments

We used OpenCap, a free and open-source software platform enabling smartphone video-based biomechanical analysis.^20^ OpenCap computes three-dimensional anatomical marker positions and skeletal joint angles from inverse kinematics. We selected the OpenCap settings that use HRNet pose estimation^24^ and a deep learning model to augment sparse 3D keypoints into a richer anatomical marker set.^25^ Within OpenCap, joint angles are computed by the OpenSim^2^ Inverse Kinematics tool and a biomechanically constrained skeletal model.^26^ All recordings were captured using either two or three cameras. A typical three-camera setup was developed (Figure S4); however, some trials were recorded with only two cameras due to space and logistical constraints (e.g., recording in a clinic hallway). In a prior validation experiment, OpenCap’s accuracy did not differ between two-and three-camera setups.^20^ All videos were recorded with at least 60 frames per second. The capability to record at 120 frames per second was developed mid-way into data collection, so some gait videos were collected at 120 frames per second to reduce the effects of motion blur during the run activity. Simultaneous with OpenCap recordings, TFTs were measured using stopwatches by evaluators trained by two physical therapists and neuromuscular disease outcomes experts in the Stanford Neuromuscular Clinic. TFTs are known to have very high inter-rater reliability with intraclass correlation above 0.99.^27,28^

### Analysis of video-based reproduction of TFTs

We computed Pearson correlations between our video-derived time scores analogous to TFTs and their corresponding human-measured TFTs. In the case of gait trials, we estimate the TFT time by dividing 10 meters by the gait speed measured in the video field of view. In the case of timed up-and-go and sit-to-stand trials, our algorithm automatically computes the time taken to complete the task. These video-based TFT surrogates do not exactly match what the human is measuring with TFT times. For walking and running, this is because the full ten meters are not visible in the camera field of view. For timed up-and-go and sit-to-stand, discrepancies arise because clinicians time when participants contact the chair seat, while our algorithm is based on center-of-mass elevation. Furthermore, all TFTs are subject to human error in measuring timing of heel strikes or chair contact. Thus, we sought to test whether video analysis can predict human-measured TFT times. To do this, we trained a linear regressor to predict human-measured TFTs from corresponding video-based TFT surrogates. We assessed prediction performance on unseen examples using leave-one-subject-out cross-validation, measuring accuracy with Pearson correlation and mean absolute error (MAE).

To measure test-retest reliability, 13 participants with DM repeated the data collection on the following day. Repeat trials were only used for test-retest analysis. Reliability was measured using intraclass correlation (ICC) with a two-way mixed effects model for absolute agreement of single measurements.^29^

### Analysis of disease classification

To study whether video-based biomechanical metrics can detect disease-specific movement patterns that TFTs do not, we compared the performance of binary classification models trained on all participant data using leave-one-subject-out cross-validation on either video-based metrics or TFT times. For participants in the test-retest experiment, only the first session was included. We used a linear support vector classifier, which is data-efficient and robust to input perturbations.^30^ Input features were standardized (zero mean, unit standard deviation) prior to model fitting. The classifier’s regularization parameter was chosen in each training fold using inner 3-fold cross validation. Model performance was quantified by balanced accuracy of predictions, which represents the percent of correct predictions, accounting for imbalance in class sizes. We compared different models’ performance using the McNemar test of proportions which considers, in the cases where the predictions of two models disagree, whether one is more often correct.^31^

### Analysis of disease-specific movement signatures

To explore which video-derived features were different between FSHD and DM, we performed two-sided Kolmogorov-Smirnov tests on all 34 engineered features. This was an exploratory analysis, and we applied Bonferroni corrections to control for type I errors due to multiple hypothesis testing.

### Analysis of feature importance

To analyze the relative importance of the video metrics for each disease, we used all video-based features to train support vector classifiers distinguishing FSHD from controls and distinguishing DM from controls. We computed Shapley additive explanations (SHAP) scores,^32^ which represent the amount each feature changes the classifier’s probability of predicting an individual as having a neuromuscular disease. We measured feature importance as the mean absolute SHAP score across all participants.

## Results

### Video-based reproduction of TFTs

Video-based TFT surrogates are strongly correlated with human-measured TFT times with Pearson r > 0.98 (Figure S7). Linear models using video-based TFT surrogates accurately predict human-measured TFT times with Pearson r > 0.98 (Figure 2a-d). To contextualize these results, we compare the MAE of video-predicted TFTs against reported distribution-based minimum clinically important difference (MCID). MCIDs for DM are available for 10-meter walk and timed up-and-go. In both cases, our MAE is below the MCID (10-meter walk: MCID = 0.69 s, MAE = 0.31 s; timed up-and-go: MCID = 0.92 s, MAE = 0.22 s).^33^ For the 10-meter run, our MAE (0.24 s) falls under the MCID for Duchenne muscular dystrophy (2.3 s).^34^ The MCID for 5-time sit-to-stand in neuromuscular disease is not available in existing literature. Video-based TFT predictions showed similar test-retest reliability as human-measured TFT times and Brooke scores. Among other video features, 70% had moderate reliability or better (Figure S1).

**Figure 2:**
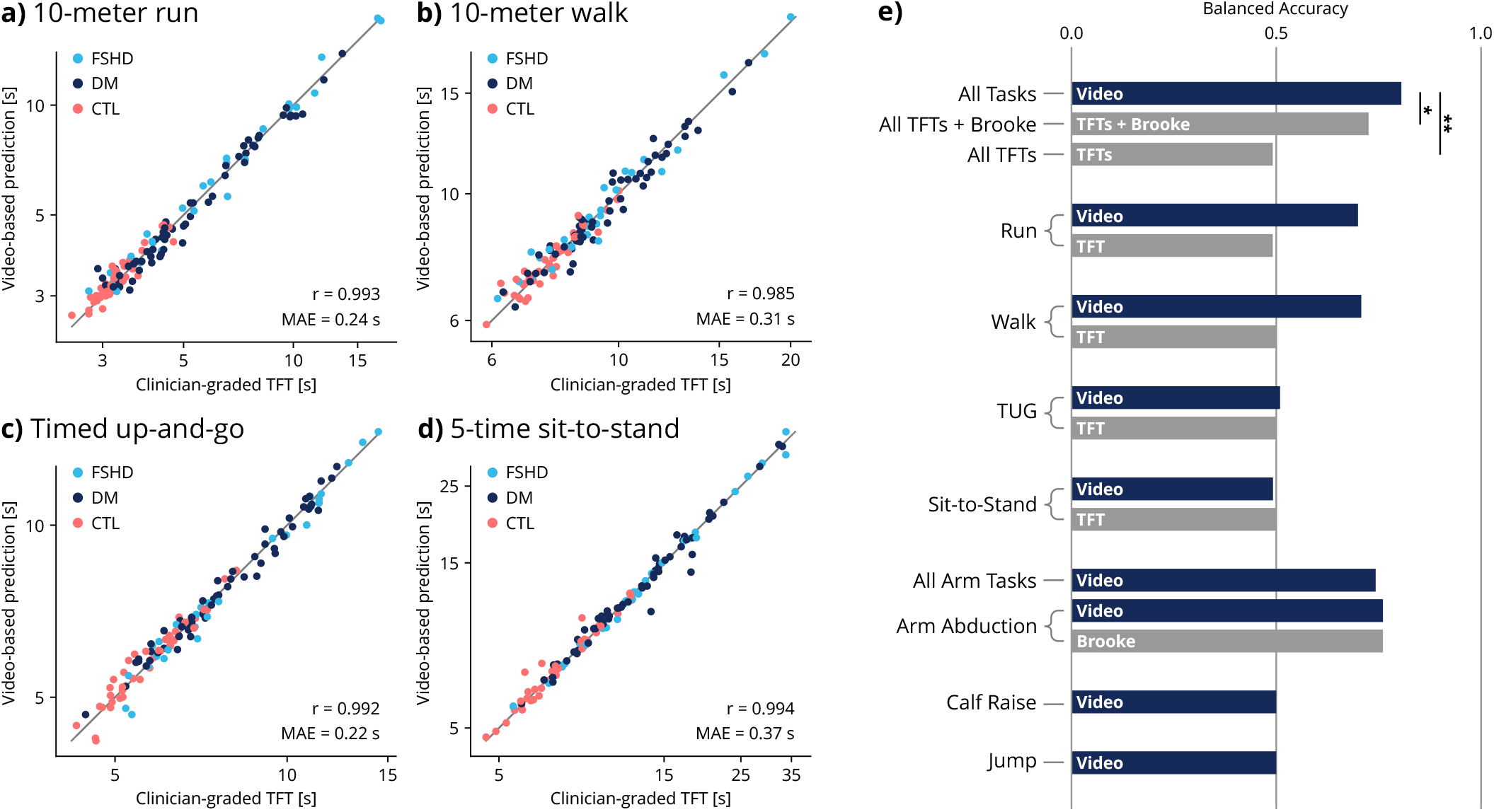
Construct and discriminant validity. To estimate human-graded timed function tests (TFT) from video-based biomechanical data, we computed related features (e.g., walk speed) and trained a linear regressor to predict the TFT time (e.g., walk time) based on the video-based features to account for systematic diZerences between the scores. Here, we show the video-predicted TFT times using a linear regressor fit with leave-one-subject-out cross validation for **a)** 10-meter run, **b)** 10-meter walk, **c)** timed up-and-go (TUG), and **d)** 5-time sit-to-stand (5xSTS) activities. **e)** Binary classification models were trained to distinguish between participants with facioscapulohumeral muscular dystrophy (FSHD) and myotonic dystrophy (DM). A model using video-based biomechanical metrics from nine activities is more accurate than models using timed function test tests (TFTs) and the Brooke Upper Extremity Scale. Video models using running and arm range of motion tasks also outperformed clinical scores. Classification model performance is measured by balanced accuracy, which corrects for unequal class sizes; 0.5 corresponds to always predicting the more common disease. Statistical diZerences in performance are tested using the McNemar test (*p < 0.05, **p < 0.01). (r = Pearson correlation, MAE = mean absolute error, CTL = control) Source: created by authors

### Disease classification

A classifier trained on video metrics derived from all nine activities performed better (balanced accuracy = 82%) than one trained on TFT times (balanced accuracy = 50%, p = 0.001) and one trained on TFT times and Brooke scores (balanced accuracy = 72%, p = 0.021) (Figure 2e). Video models using features from individual activities performed as well or better than TFT models across all activities. TFT times from individual tasks classify diseases no better than random (balanced accuracy = 50%); however, video-based models classify diseases better than random using running features (balanced accuracy = 68%) or walking features (balanced accuracy = 66%). This is because video-based biomechanical analysis captures some disease-specific gait parameters that are not reflected in TFT times. For example, video-based biomechanical analysis detected differences in running stride length between FSHD and DM participants (p = 0.044), while running speed (analogous to the TFT time) was not significantly different (Figure S2).

### Disease-specific feature importance

The most important features for detecting FSHD are upper-limb activities representing shoulder and elbow range of motion (Figure 4a). By contrast, the most important features for detecting DM are from lower-limb activities such as running, walking, and jumping (Figure 4b). These observations are consistent with the known early clinical presentations of upper-limb and shoulder girdle weakness in FSHD and distal lower-extremity weakness in DM.

### Disease-specific movement signatures

Compared to the DM group, the FSHD group had shorter stride lengths (p = 0.037) and higher ankle heights (p = 0.008) during the swing phase of walking (Figure 3a). This is consistent with the presentation of foot drop in FSHD due to ankle dorsiflexor weakness.^9^ There was no significant difference in walking speed between FSHD and DM, consistent with the inability of the walking TFT model to classify diseases (Figure 2e).

**Figure 3:**
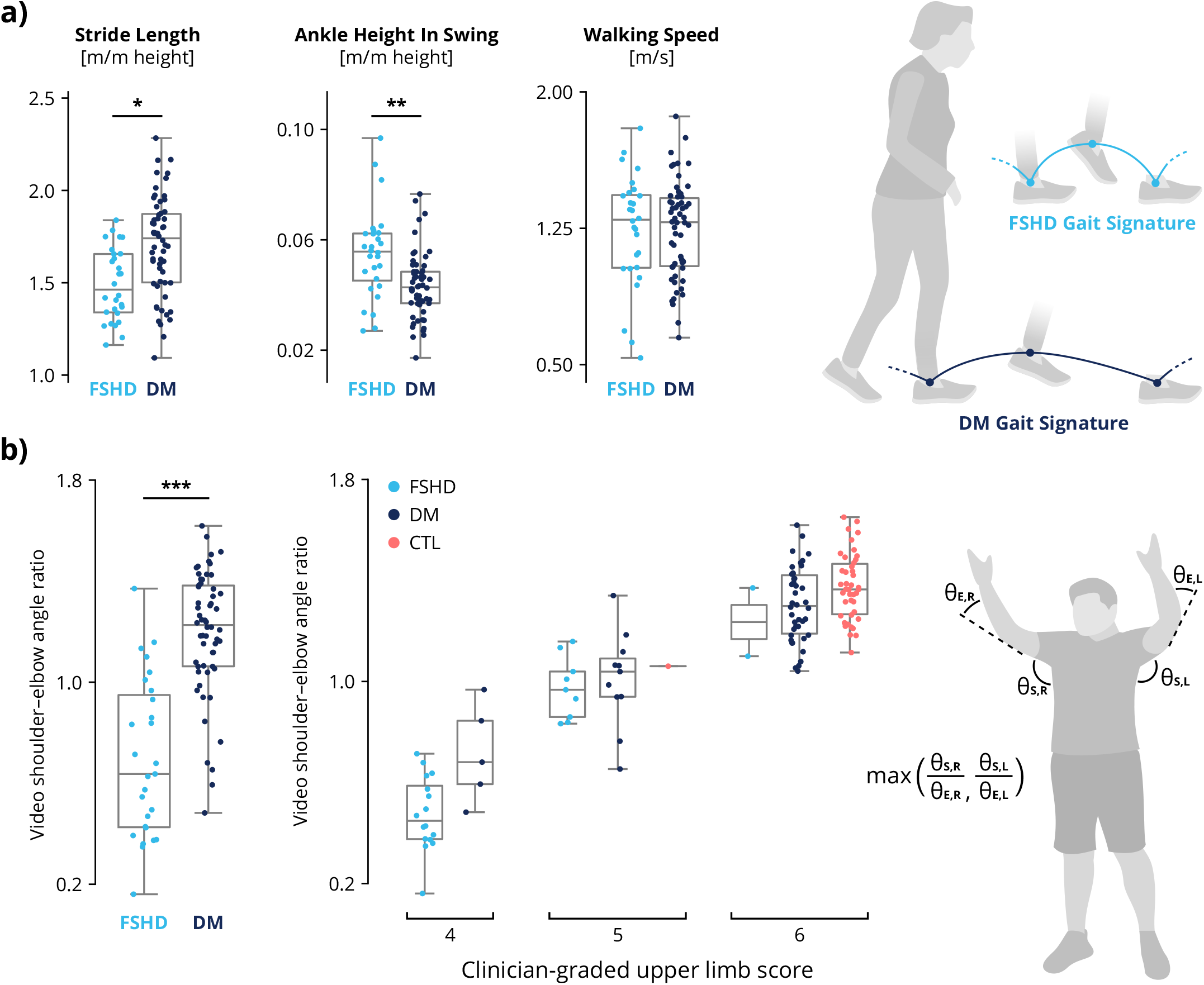
Disease-specific movement signatures. Video-based analysis reveals disease-specific movement signatures in gait and upper-limb range-of-motion. **a)** Patterns in walking gait diZer between FSHD and DM. Despite similar walking speeds, the FSHD group walked with shorter stride lengths (p = 0.037) and higher ankle height during swing (p = 0.008) than did the DM group. (“m/m height” indicates units of meters normalized by participant height in meters, i.e. dimensionless.) **b)** Video-based biomechanical analysis of upper-body function shows diZerences between diseases (p = < 0.001). A video-based metric agrees with (Spearman’s rank correlation r = 0.80). the human-graded Brooke Upper Extremity Scale, which ordinally scores the ability to raise arms overhead with straight elbows. To compute our analogous video-based metric, we calculate the ratio of shoulder abduction angle (***θ_S_***) to elbow flexion angle (***θ_E_***) and take the maximum ratio between right and left sides. Statistical significance is measured with a two-sided Kolmogorov–Smirnov test with Bonferroni corrections to control for Type I error across all 34 features. (FSHD = facioscapulohumeral muscular dystrophy, DM = myotonic dystrophy, CTL = control) Source: created by authors

Compared with DM, FSHD more strongly affects the proximal arm muscles, reducing shoulder and scapulothoracic function. We found significant differences in video-based biomechanical metrics of upper-limb tasks between diseases (Figure 3b). Ninety participants received a Brooke score of 6, indicating an ability to abduct hands overhead without elbow flexion; 23 received a score of 5 (able to raise hands overhead with some elbow flexion), and 21 received a score of 4 (unable to raise hands overhead). Our video metric comprising a composite of shoulder and elbow angles (Figure 3b) correlates with human-graded Brooke scores (Spearman’s rank correlation r = 0.80). Despite this correlation, there is high variance in the video metric among individuals with the same Brooke score, highlighting the insensitivity of ordinal scores like Brooke. Additionally, we observed significant differences between diseases in other video-based features from upper-body tasks (e.g., reachable area and isolated elbow flexion angle), showing how these metrics can capture disease-specific patterns in upper-body movement (Figure S2). Together, these results highlight the strength of a continuous video-based metric to characterize upper-limb function compared to a human-graded ordinal score.

## Discussion

Video-based 3D biomechanical analysis using OpenCap reproduces clinician-graded assessments of mobility and function while also encoding additional disease-specific insights. OpenCap video-based assessments accurately predicted TFT scores from videos (Figure 2a-d) with similar test-retest reliability (Figure S1). Unlike TFTs, video-based analysis differentiated the movement patterns of FSHD versus those of DM (Figure 2e) and identified disease-specific biomechanical impairments (Figure 3). These findings demonstrate the potential of using OpenCap-based biomechanical analysis to create functional outcome measures for neuromuscular diseases that are more sensitive than commonly used TFTs.

Video-based analysis is sensitive to disease-specific motor impairments caused by dysfunction in distinct muscle groups. For example, video-based analysis of running gait distinguishes between diseases while TFT times alone do not (Figure 2e). Video-based analysis also reveals specific gait parameter differences — such as step length and ankle height — between FSHD and DM (Figure 3a), indicating that video metrics capture salient aspects of disease signatures that TFT times do not. Furthermore, video-based metrics distinguish differences in upper-limb range of motion between FSHD and DM and capture functional variability among individuals assigned the same human-graded Brooke score (Figure 3b). These findings suggest that video-based analysis enables more sensitive upper-limb functional assessments.

One reason video features outperform TFT times for classifying diseases is that they include measures of upper-limb mobility such as shoulder and elbow range of motion. Analysis of feature importance reveals that, unlike DM, the FSHD group could be distinguished from the control group mainly in upper limb features (Figure 4). This is consistent with the known clinical presentation: the scapular stabilizers, which help raise the arms above the head, are among the most affected muscles in FSHD^10^ whereas they are less affected in DM.^11^ However, even when Brooke scores are combined with TFTs, video-based analysis of all tasks still achieves significantly better classification (Figure 2e). Thus, the advantage of video-based analysis goes beyond simply including upper-limb activities.

**Figure 4:**
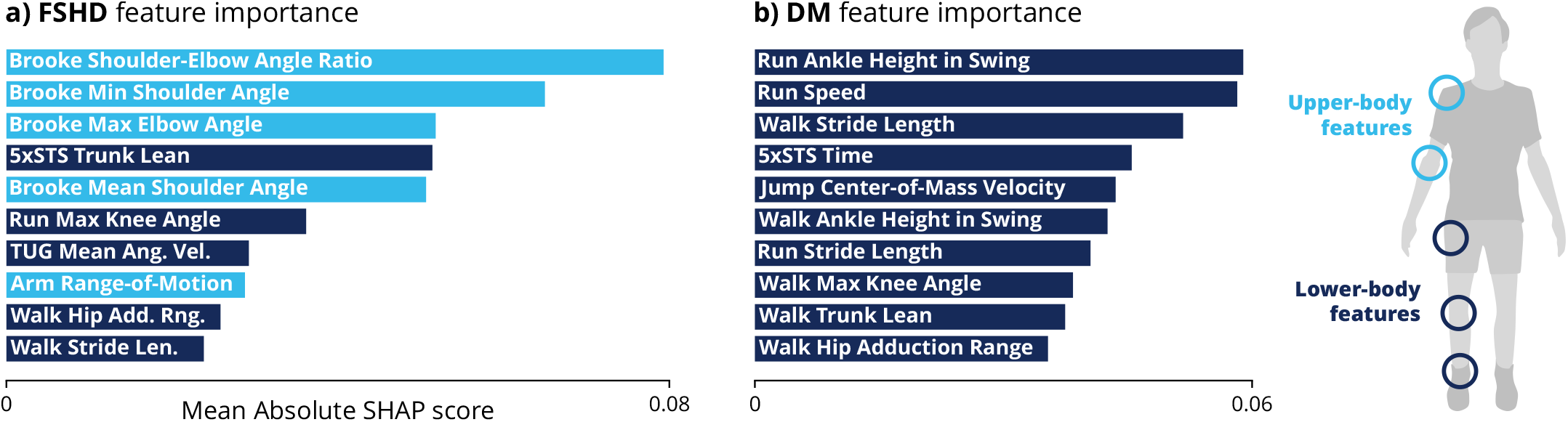
Disease-specific feature importance. Binary classification models were trained to distinguish control participants versus those with **a)** facioscapulohumeral dystrophy (FSHD) or **b)** myotonic dystrophy (DM). Feature importance is measured by the mean absolute Shapley additive explanations (SHAP) values, which indicate the average amount each feature impacts the classifier’s probability of predicting disease. The top ten features are shown here; all feature importance scores are in Figure S3. The classification models are linear support vector machines. The FSHD model relies mainly on upper-limb features, while the DM model relies mainly on lower-limb features. Descriptions of video-based features are in Table S3. (5xSTS = 5-time sit-to-stand, TUG = timed up-and-go) Source: created by authors

Our set of video-based features are not comprehensive. Different features may demonstrate different disease classification performance. For example, our features extracted from the timed up-and-go features exclude gait parameters to avoid redundancy with the walk and run tasks. An alternative to our manual feature engineering process based on pathophysiological, clinical, and biomechanical domain-knowledge is to derive data-driven features using supervised or unsupervised dimensionality reduction.^35^ It is possible that these methods could improve classification performance, but the features would be less interpretable than hand-engineered features.

The ultimate goal is to create digital outcome measures that are more sensitive and reliable than traditional TFTs. Video-based biomechanical analysis shows promise for achieving greater sensitivity than existing clinical outcome metrics.^6,7^ Some video-based biomechanical metrics have low reliability scores among test-retest participants. With the present dataset, we cannot determine what proportion of variance is attributable to day-to-day variability in task performance versus measurement error. Unsurprisingly, for both the TFT and video measures, the maximal effort tasks (e.g., run) were more repeatable than sub-maximal tasks (e.g., walk). Greater variance in an outcome measure decreases sensitivity. Future studies could choose to focus on the metrics that we found to have high reliability.

The advantages of novel digital health metrics must be balanced with the burden of data collection and analysis. In this study, the median time to record nine activities was 16 minutes, with kinematics calculations automated on OpenCap’s cloud servers. For comparison, collecting the same data using typical laboratory-based motion capture would take three or more hours including calibration, equipment setup, marker placement, activity recording, and data post-processing. OpenCap achieves near lab quality kinematics with a much lower time burden.^20^ This is especially significant for studying rare diseases. For example, our cohort of 58 individuals with DM is nearly four times larger than the largest existing study of 3D kinematics in DM.^36^ By collecting data in five locations, including national conferences, we enrolled participants from 14 different states within the United States. Compared with traditional motion capture, OpenCap is inexpensive to scale. Our high-throughput data collection included six simultaneous recording stations. Taken together, the speed, portability, and scalability of OpenCap enabled a cohort seven times larger (n=129) than the average laboratory-based biomechanics study (n=18).^37,38^

It is important to acknowledge the limitations of this study. First, further research is required to determine if the advantages of video-based analysis generalize beyond FSHD and DM to other neuromuscular diseases. However, our findings imply that in these conditions, OpenCap video-based biomechanical analysis provided more information than a commonly used functional outcome metric and has the potential to improve the creation of movement biomarkers and functional outcome metrics for other conditions. It is also important to consider how feature engineering can impact the generalizability of classification results. During feature engineering the research team was not blinded to all participant diagnoses since the research team also performed data collection. However, we believe this poses minimal risk of overfitting because clinicians viewed less than 30 of the 1,290 OpenCap recordings in our dataset during feature engineering. Second, our convenience sample includes a low proportion of female participants with FSHD because FSHD presents earlier and more severely in men (Table S1). The low number of females with FSHD impacts disease classification on this group (Table S4). However, a video-based classifier trained and tested on only male participants does not have worse disease classification (balanced accuracy = 83%), suggesting that our classifier does not depend on sex-linked movement patterns to differentiate diseases. Third, while video-based analysis was faster, cheaper, and easier than traditional methods, our multi-camera setup complicates at-home and continuous measurements. Wearable sensors could better capture long-term patterns in movement and behavior,^6,39–42^ but video is better suited for periodic, high-resolution assessments of whole-body biomechanics in controlled settings. Single-camera biomechanical analysis will likely become possible in the near future,^43–45^ which may enable more frequent at-home measurements or even opportunistic sensing of uninstructed activities of daily living. Finally, the present analysis is a cross-sectional assessment of construct and discriminant validity. Further work should determine repeatability, correlation with functional and patient-reported outcomes, minimum clinically important differences, and sensitivity to natural progression and therapeutic interventions.

This work shows that OpenCap video-based biomechanical analysis can both reproduce TFTs and measure patterns of movement in FSHD and DM that TFTs cannot. The future clinical implications of video-based functional assessments lie in their ability to enable more sensitive evaluation and monitoring of motor function, with applications in improving diagnosis, accelerating treatment development, and enabling personalized patient management.

## Data sharing

The datasets generated and analyzed for this study are freely available in the “Kinematics and timed function tests of facioscapulohumeral muscular dystrophy and myotonic dystrophy” data repository, available at https://doi.org/10.5281/zenodo.13788592. The code for feature extraction, figure generation, and statistics is available in the following GitHub repository: https://github.com/stanfordnmbl/opencap-fshd-dm-analysis.

## Acknowledgements

We thank the 42 volunteers who helped facilitate recruitment, data collection, and data curation, especially Lin Karman, Sarah Ismail, Paxton Ataide, and Carmichael Ong. We are grateful to Trevor Hastie for consultation on statistical methods. Thank you to the Myotonic Dystrophy Foundation, and the FSHD Society for their support of our study. We especially wish to thank the 129 study participants without whom this research would not have been possible. This study was funded by the Wu Tsai Human Performance Alliance, Mobilize Center under NIH Grant P41EB027060, and the Myotonic Dystrophy Foundation Early Career Research Grant. The funders played no role in study design, data collection, analysis and interpretation of data, or the writing of this manuscript.

## Author contributions

P.S.R., S.D.U., C. M., J.D., T.D., and S.L.D. conceptualized and designed the study.

S.D.U., C.M., A.F., J.M., and T.D. developed the methods used to perform the experiments.

P.S.R., S.D.U., C.M., A.F., J.M., S.C., S.V., and T.D. conducted the experiments.

P.S.R. synthesized results and figures.

P.S.R. and S.D.U. drafted the manuscript.

C.M., A.F., J.M., S.C., S.V. J.D., T.D., and S.L.D. revised the manuscript.

