## Supplement for "Video-based biomechanical analysis captures disease-specific movement signatures of different neuromuscular diseases"

**Supplementary Table 1: Detailed study participant characteristics.** Note that some individuals participated at multiple locations. The smaller proportion of female FSHD participants is due to the population trends since FSHD is more commonly asymptomatic in females than males.[1]
(FSHD = facioscapulohumeral muscular dystrophy, DM = myotonic dystrophy, TYP = typical movement)

| | **Total** | **FSHD** | **DM** | **Typical** |
|---|---|---|---|---|
| Participants | 129 | 28 | 58 | 43 |
| **Location** | | | | |
| High-throughput study | 94 | 26 | 27 | 41 |
| Community events | 30 | 1 | 27 | 2 |
| Neuromuscular clinic | 5 | 1 | 4 | 0 |
| **Sex** | | | | |
| Female (%) | 57 (44%) | 4 (14%) | 31 (53%) | 22 (51%) |
| **Age Group** | | | | |
| 10–19 | 4 | 0 | 2 | 2 |
| 20–29 | 22 | 4 | 10 | 8 |
| 30–39 | 32 | 6 | 16 | 10 |
| 40–49 | 25 | 3 | 16 | 6 |
| 50–59 | 23 | 7 | 7 | 9 |
| 60–69 | 16 | 6 | 4 | 6 |
| 70–79 | 5 | 1 | 3 | 1 |
| 80–89 | 2 | 1 | 0 | 1 |
| Mean (SD) | 43 (16) | 48 (16) | 41 (14) | 44 (17) |
| **BMI** | | | | |
| < 18.5 | 8 | 1 | 6 | 1 |
| 18.5–25 | 65 | 13 | 27 | 25 |
| 25–30 | 33 | 10 | 11 | 12 |
| > 30 | 23 | 4 | 14 | 5 |
| Mean (SD) | 25.2 (5.5) | 25.8 (4.6) | 25.4 (6.6) | 24.7 (4.4) |

**Supplementary Table 2: Physical activities.** (RoM = range of motion, TFT = timed function test)

| Activity | Description | Clinical Assessment |
|---|---|---|
| Run | As fast as able, run or walk 10 meters from standing cold start | 10m walk/run test (TFT) |
| Walk | At self-selected comfortable pace, walk 10 meters from standing cold start | 10m walk test (TFT) |
| TUG | As fast as able, stand up, walk 3 meters, turn around a cone, walk back, and sit down | Timed up-and-go (TFT) |
| 5xSTS | As fast as able, rise from chair and sit down again five times with arms crossed over chest | 5-time sit-to-stand (TFT) |
| Brooke | Abduct arms overhead with straight elbows | Brooke Upper Extremity Scale |
| ArmRoM | Raise arms with straight elbows as high as able in four directions (0º flexion, 45º scaption, 90º abduction, 180º extension); horizontally sweep straight arms at waist and eye levels | N/A |
| Curls | Flex elbows as far as able, starting from full extension | N/A |
| ToeStand | Rise to toes for 30s or as long as able | N/A |
| Jump | Jump as high as able; if unable, rise from a squat as fast as able | N/A |

**Supplementary Table 3: Descriptions of engineered features.** "m/m" means meters divided by participant height, i.e. dimensionless. "$m^2/m^2$" means area divided by participant height squared, i.e. dimensionless. "$ms^{-1}/m$" means speed normalized by participant height, i.e. units of 1/s.

| Activity | Feature | Description | Units |
|---|---|---|---|
| Run | Speed | Mean speed | m/s |
| Run | Stride Time | Mean stride time | s |
| Run | Stride Length | Median stride length; height normalized | m/m |
| Run | Ankle Height in Swing | Median ankle height in swing; height normalized | m/m |
| Run | Center-of-Mass Sway | Center of mass lateral velocity standard deviation | m/s |
| Run | Trunk Lean | Mean absolute lateral trunk angle | ° |
| Run | Hip Abduction Range | Hip abduction/adduction angle range; L/R averaged | ° |
| Run | Max Knee Angle | Max knee angle; L/R averaged | ° |
| Walk | Speed | Mean speed | m/s |
| Walk | Stride Time | Mean stride time | s |
| Walk | Stride Length | Median stride length; height normalized | m/m |
| Walk | Ankle Height in Swing | Median ankle height in swing; height normalized | m/m |
| Walk | Sway Velocity | Center of mass lateral velocity standard deviation | m/s |
| Walk | Trunk Lean | Mean absolute lateral trunk angle | ° |
| Walk | Hip Abduction Range | Hip abduction/adduction angle range; L/R averaged | ° |
| Walk | Max Knee Angle | Max knee angle; L/R averaged | ° |
| TUG | Time | Time from rise to sit | s |
| TUG | Mean Angular Velocity | Mean angular velocity during turn | °/s |
| TUG | Max Angular Velocity | Max angular velocity during turn | °/s |
| 5xSTS | Time | Time to complete 5 repetitions | s |
| 5xSTS | Trunk Lean | Maximum trunk angle | ° |
| 5xSTS | Stance Width | Max distance between ankles; height normalized | m/m |
| Brooke | Mean Shoulder Angle | Maximum of L/R mean shoulder angle | ° |
| Brooke | Min Shoulder Angle | Maximum of L/R min shoulder angle | ° |
| Brooke | Max Elbow Angle | L/R max elbow angle when arms overhead | ° |
| Brooke | Shoulder-Elbow Angle Ratio | Max ratio of L/R mean shoulder angle to L/R mean elbow angle | °/° |
| ArmRoM | Reachable Area | Convex hull area of wrist trajectories; normalized by height squared | $m^2/m^2$ |
| Curls | Min Elbow Angle | L/R min of elbow flexion angle | ° |
| Curls | Mean Elbow Angle | L/R mean of elbow flexion angle | ° |
| ToeStand | Center-of-Mass Height | Integral of center-of-mass elevation; height normalized | $ms^{-1}/m$ |
| ToeStand | Heel Height | Integral of mean heel elevation; height normalized | $ms^{-1}/m$ |
| ToeStand | Trunk Lean | Integral of trunk angle | °/s |
| ToeStand | Ankle Angle | Integral of ankle angle | °/s |
| Jump | Center-of-Mass Velocity | Max center-of-mass velocity | m/s |

**Supplementary Table 4: Classifier performance by sex.** To consider effects of sex imbalance on classifier accuracy, we show a breakdown of the proportion of male and female participants with FSHD and DM who are correctly and incorrectly classified by the all-task video based feature model (Fig. 2b).

| | **Has FSHD** | **Has DM** | |
|---|---|---|---|
| **Predicted FSHD** | 1 F, 19 M<br>5% F | 2 F, 2 M<br>50% F | 3 F, 21 M<br>12% F |
| **Predicted DM** | 3 F, 5 M<br>38% F | 29 F, 25 M<br>54% F | 32 F, 30 M<br>52% F |
| | 4 F, 14 M<br>14% F | 31 F, 27 M<br>53% F | |

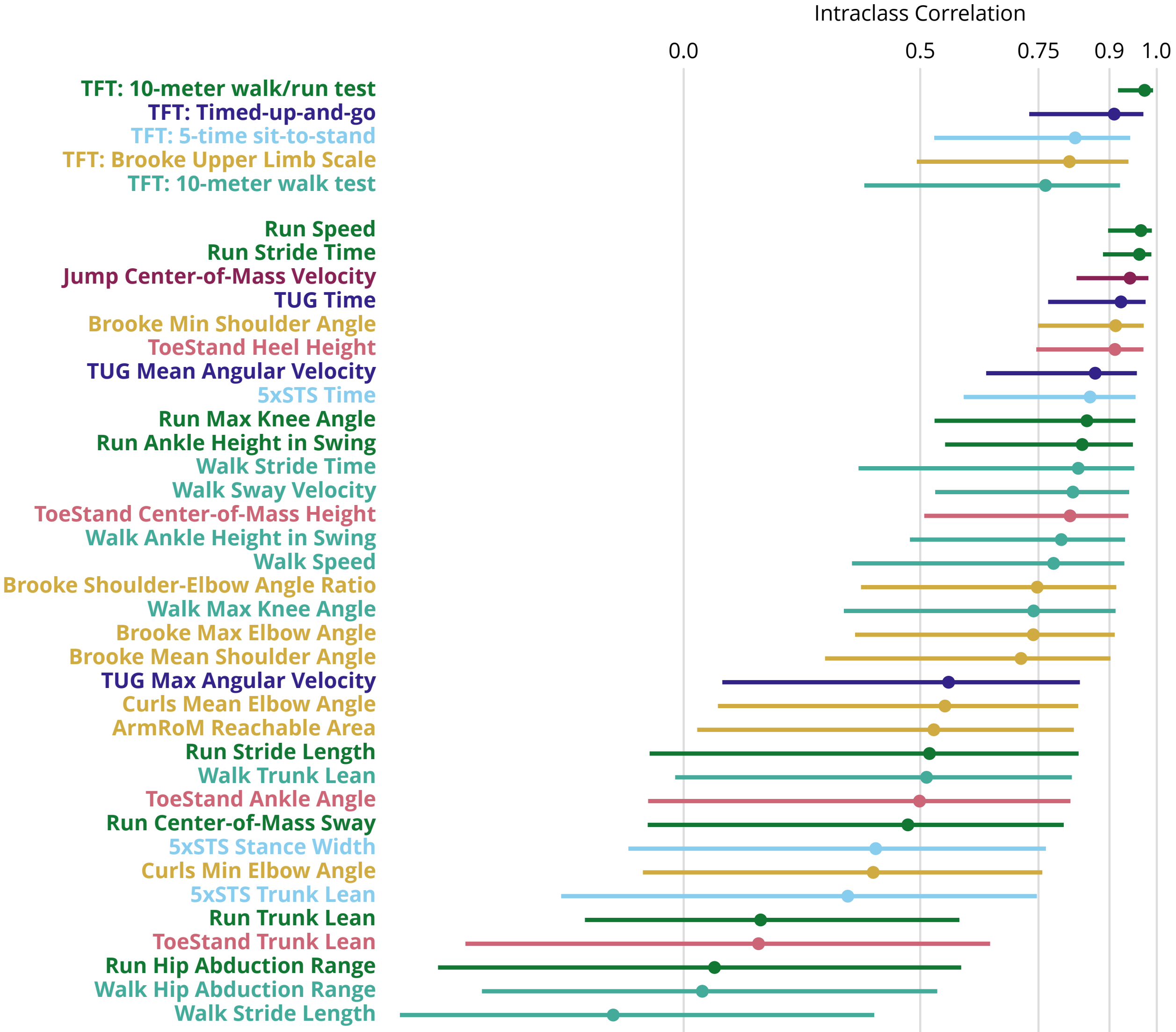


**Supplementary Fig. 1: Test-retest reliability.** The reliability of video-based metrics, timed function tests (TFTs), and Brooke Upper Limb Scale are measured by intraclass correlation (ICC). Bars indicate 95% confidence intervals. ICC > 0.5 indicates "moderate" reliability, ICC > 0.75 indicates "good" reliability, and ICC > 0.9 indicates "excellent" reliability. (5xSTS = 5-time sit-to-stand, TUG = timed up-and-go). Colors denote features from the same activity.

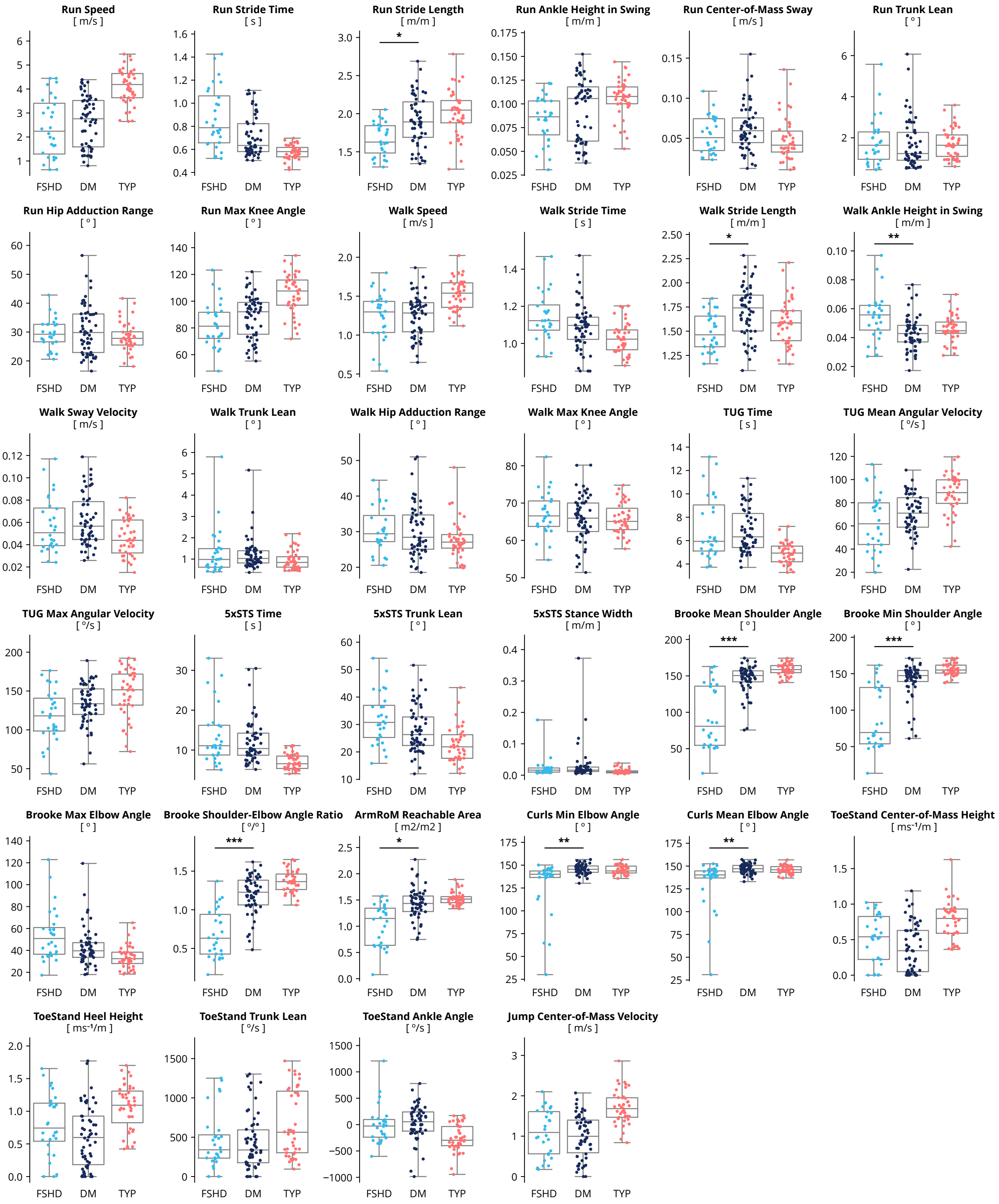


**Supplementary Fig. 2: Disease differences by feature.** For all features, we compared FSHD and DM groups using a two-sided Kolmogorov-Smirnov tests with Bonferroni corrections for multiple hypothesis testing (*p < 0.05, **p < 0.01, ***p < 0.001). "m/m" means meters divided by participant height, i.e. dimensionless. "$m^2/m^2$" means area divided by participant height squared, i.e. dimensionless. CTL groups were not statistically compared to other groups. "$ms^{-1}/m$" means speed normalized by participant height, i.e. units of 1/s. (FSHD = facioscapulohumeral muscular dystrophy, DM = myotonic dystrophy, CTL = typical movement)

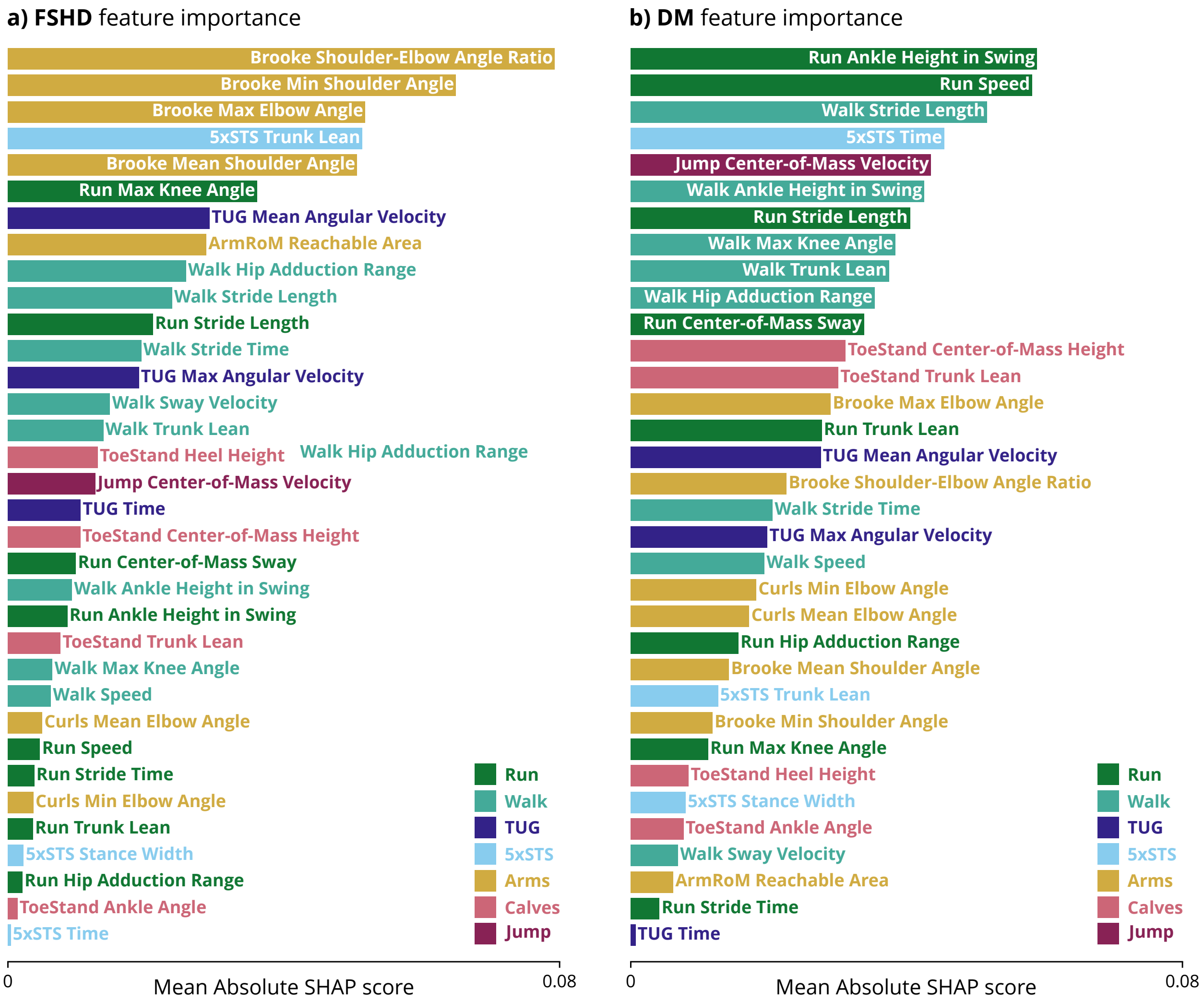


**Supplementary Fig. 3: Feature importance.** Binary classifiers were trained to distinguish control participants versus those with **a)** facioscapulohumeral dystrophy (FSHD) or **b)** myotonic dystrophy (DM). The classifiers are support vector machines with radial basis function kernels. SHapley Additive exPlanations (SHAP) scores describe the contribution of each video-derived feature towards the probability of neuromuscular disease for a given training example. For each feature, we plot the mean absolute SHAP score across all training examples. Units are in terms of probability (e.g. a mean absolute SHAP score of 0.05 means the feature impacts the predicted probability by 5% on average). (5xSTS = 5-time sit-to-stand, RoM = range of motion, TUG = timed up-and-go)

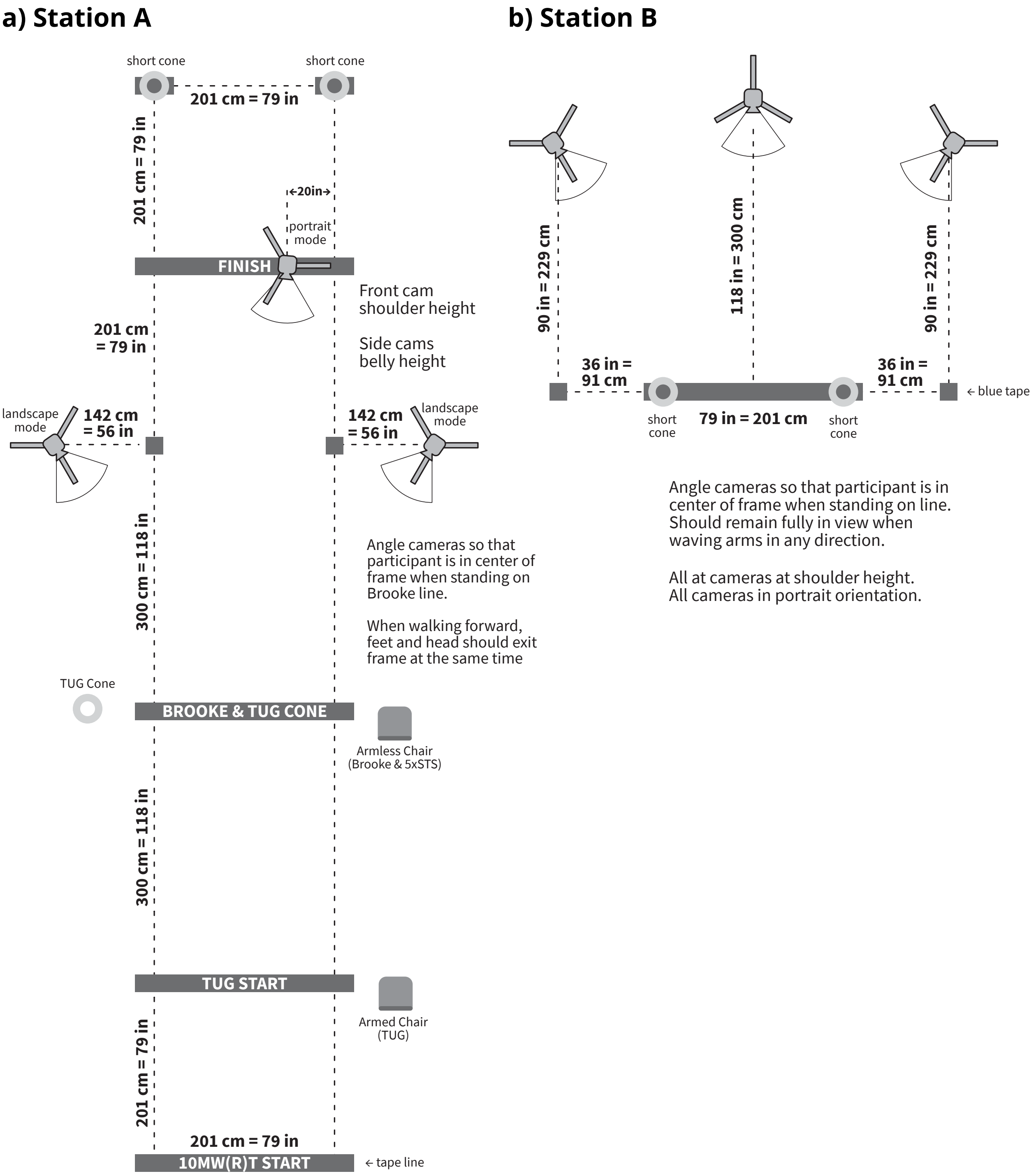


**Supplementary Fig. 4: Experimental setup.** Printable versions available upon request. Activities performed at each station are listed in Supplementary Fig. 5 and Supplementary Fig. 6. "10MW(R)T START" marks the start of the 10-meter walk and 10-meter run tests. TUG = timed up-and-go. Brooke is a seated upper body assessment. 5xSTS = five-time sit-to-stand.

pID

### STATION **A**

Clinician Initials

**Neutral**

**Stand straight, feet hip-width apart, toes forward, arms slightly out to the sides.**
[Demonstrate]
**Stay still until we tell you to relax.**

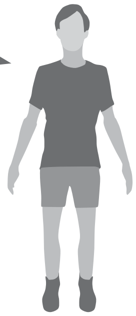

**Brooke**

**First I will demonstrate the movement and then we will do it together. Start with hands at your side, then raise them over your head while keeping your elbows straight. Try to keep your arms behind your ears as you bring them over your head.**
[Demonstrate slow straight arm abduction, thumbs up, touch palms together overhead]
**When I say go we will perform this motion together.**
[Start OpenCap recording]
**Ready, set, follow me...** [Perform activity with participant.]
Assign Brooke Score:

- If completed task with no compensations → **6**
- If hands above head, but only with compensation (e.g. elbows bent, shoulders shrugged, head tilted forward, or arms in front of body) → **5**
- If hands not above head, give participant cup with 200g weight
  - If able to lift weighted cup from lap to chin (both hands okay) → **4**
  - If unable, remove 200g weight
    - If able to lift empty cup from lap to chin (both hands okay) → **3**
    - If unable → **2**

**10MWT**

s

**This is a 10 meter walk test. Stand with your feet behind the line. First I will ask you to raise your hand like this. Then return your hands to your sides. Then I say go, I want you to walk at your comfortable walking speed past the cones.**
[Demonstrate]
**We won't be encouraging you during the test, but we want you to walk at your normal speed. Please don't talk while you're walking.**
[Start OpenCap recording] [Cue hand raise]
**Ready, set, go!** [Start timer on "go"] [Stop timer on first heel strike fully past 10m tape mark]

**10MWRT**

s

**Next we'll test how fast you can move. Start with your feet behind the line. First I will ask you to raise your hand. Then return your hands to your sides. Then, when I say go, I want you to move as fast as you can safely past the cones. If you are able to safely run, you may do so; if not, walk as fast as you can. Remember to go as fast as you can past the cones.**
**We want you to give your best effort.**
[Start OpenCap recording] [Cue hand raise]
**Ready, set, go!** [Start timer on "go"] [Stop timer on first heel strike fully past 10m tape mark]

**TUG**

s

**This is a timed up-and-go test. Sit as far back in the chair as possible with your feet flat on the floor. Start with your arms on the armrests. First I will ask you to raise your hand. When I say go, stand up, then walk as fast as you can around the cone, return to the chair and sit down.**
[Demonstrate]
**Try not to use your arms to push off unless you have to. You can walk fast, but do not run or skip.**
[Start OpenCap recording] [Cue hand raise]
**Ready, set, go!** [Start timer on "go"] [Stop timer when bottom touches chair]

**5xSTS**

s

**This is a five-time-sit-to-stand test. Sit as far back in the chair as possible with your feet flat on the floor. Keep your arms crossed over your chest.**
[If unable to cross hands over chest, can cross over stomach]
**First I will ask you to raise your hand. Then cross your arms again. Then when I say go, I want you to stand up and sit back down five times as quickly as possible. I will count out loud while you do these. Make sure you come to a full upright standing position and touch your bottom to the chair every time. Go as fast as you can without using your arms.**
[Note: if participant needs to spread feet to stand, must return under base of support before sitting.]
[Start OpenCap recording] [Cue hand raise]
**Ready, set, go!** [Start timer on "go"] [Stop timer when bottom touches chair]

**Supplementary Fig. 5: Experimenter script A.** Printable version available upon request.

### STATION **B**

**Neutral**

**Stand straight, feet hip-width apart, toes forward, arms slightly out to the sides.**
[Demonstrate]
**Stay still until we tell you to relax.**

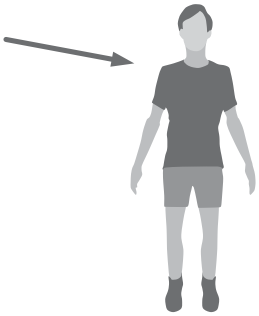

**ArmRoM**

**For this task you will move your arms in 6 directions.**
**First I will show you the movements and then we will do them together.**
Demonstrate movements:
- Arm raise: flexion (0º from AP axis)
- Arm raise: scaption (45º from AP axis)
- Arm raise: abduction (90º from AP axis)
- Arm raise: extension (180º from AP axis)
- Horizontal sweep: waist height ("hula hoop")
- Horizontal sweep: head height, elbow at eye level (or as high as able)

**When I say go we will perform those motions together.**
**Each time I want you to move your arms in a smooth controlled way.**
**I want you to move through your full range of motion. Try not to bend your elbows as much as you can.**
[Start OpenCap recording] [Note: no hand raise]
**Ready, set, follow me...** [Perform activity with participant.]

**Curls**

**Now we will do a series of elbow and wrist movements. First I will show you the movements and then we will do them together.**
Demonstrate movements:
- Elbow RoM: full extension (arms at sides, palms forward) → full flexion (touch shoulders) → full extension
- Elbows at side, 90º flexion (or as high as able), keep elbows still while doing:
  - Supinated wrist RoM (palms up): full extension (fingers down) → full flexion (fingers up) → full extension
  - Pronated wrist RoM (palms down): full flexion (fingers down) → full extension (fingers up) → full flexion

**When I say go we will perform those motions together. When you start doing the wrist motions try and keep your elbow at 90º. Each time, try and move your arms in a smooth controlled way.**
[Start OpenCap recording] [Note: no hand raise]
**Ready, set, follow me...** [Perform activity with participant.]

**ToeStand**

**First I will ask you to raise your hand. Then return your hands to your sides. Then when I say go I want you to rise onto your tip toes as quickly as possible and try and hold it there for 30 seconds. I want you to be safe. Try to keep your balance for as long as possible. If you drop down, try to rise up again until I say stop. Try to stand as stable as possible. If you need to, you may move your feet to stay upright.**
[Start OpenCap recording] [Cue hand raise]
**Ready, set, go!** [Start timer on "go"] [Cue 15 sec mark, count down last 5 seconds]

**Jump**

**First I will ask you to raise your hand. Then when I say go, I want you to try and jump as high as you can.**
[Demonstrate with countermotion]
**I want you to be safe so only do what you feel like you are able to do.**
**If you are unable to jump, simulate a jump by bending your knees and standing upright as fast as you can.**
[Start OpenCap recording] [Cue hand raise]
**Ready, set, go!**

**NOTES**

**Supplementary Fig. 6: Experimenter script B.** Printable version available upon request.

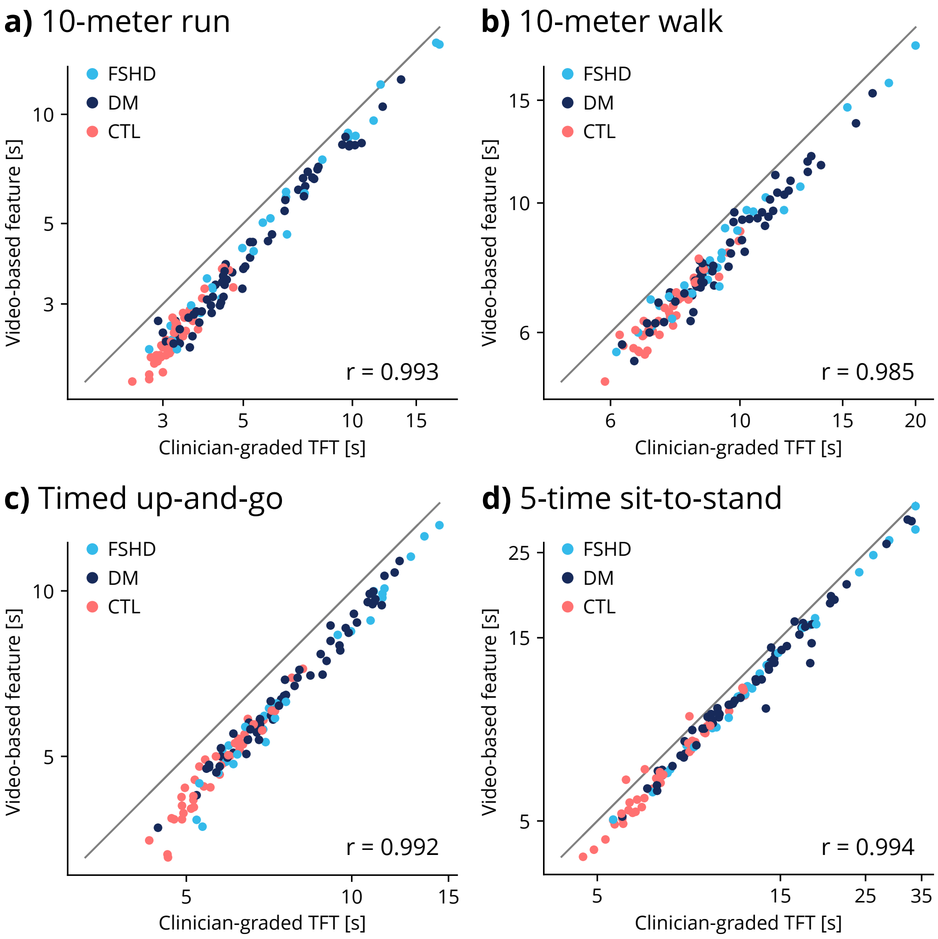


**Supplementary Fig. 7: Correlation of TFTs versus video-derived TFT surrogates.** We computed the correlation of human-measured timed function tests (TFTs) versus video-based metrics analogous to TFTs, for all all four TFTs: **a)** 10-meter run, **b)** 10-meter walk, **c)** timed up-and-go (TUG), and **d)** 5-time sit-to-stand (5xSTS) activities. (r = Pearson correlation, FSHD = facioscapulohumeral muscular dystrophy, DM = myotonic dystrophy, CTL = control)

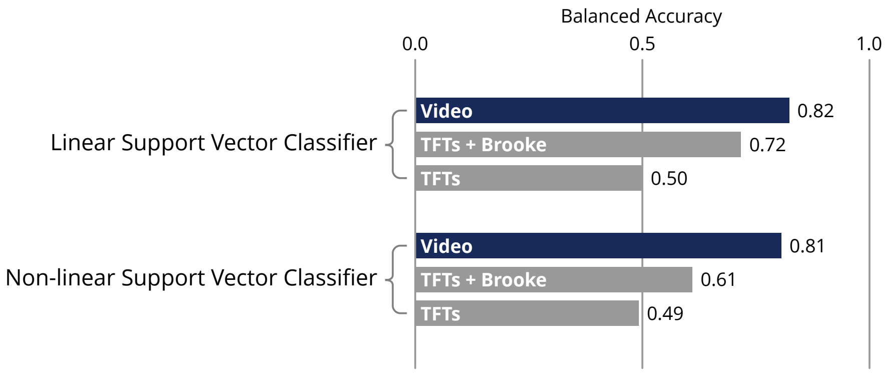


**Supplementary Fig. 8: Comparison of linear and non-linear models.** We compared the performance of support vector classifiers with linear and non-linear kernels. The non-linear kernel is a radial basis function. Linear models achieve a higher balanced accuracy for disease classification using video features, timed function test (TFT) times, and Brooke Upper Extremity Scale scores.
